# Dose and duration of IFNγ pre-licensing interact with donor characteristics to influence the expression and function of IDO in MSCs

**DOI:** 10.1101/855270

**Authors:** Devlin Boyt, Lauren Boland, Anthony J. Burand, Alex Brown, James A. Ankrum

## Abstract

Human mesenchymal stromal cells (MSCs) are a leading cell therapy candidate for the treatment of immune and inflammatory diseases due to their potent regulation of immune cells. MSC expression of indoleamine-2,3-dioxygenase (IDO) upon interferon gamma exposure has been proposed as both a sentinel marker and key mediator of MSC immunomodulatory potency. Rather than wait for *in vivo* exposure to cytokines, MSCs can be pre-licensed during manufacturing to enhance IDO expression. In this study, we systematically examine the relative role the dose of interferon gamma, the duration of pre-licensing, and the donor of origin plays in dictating MSC production of functional IDO. We find that across three human MSC donors, MSCs increase their expression of IDO in response to both increased dose of interferon gamma and duration of pre-licensing. However, with extended pre-licensing, the expression of IDO no longer predicts MSCs ability to suppress activated peripheral blood mononuclear cells. In addition, pre-licensing dose and duration are revealed to be minor modifiers of MSCs inherent potency, and thus cannot be manipulated to boost poor donors to the levels of high-performing donors. Thus, the dose and duration of pre-licensing should be tailored to optimize performance of specific donors and an emphasis on donor selection is needed to realize significant benefits of pre-licensing.

## Introduction

The immunomodulatory phenotype of mesenchymal stromal cells (MSC) has been applied in clinical trials with notable successes in the treatment of graft versus host disease^1,2^ and Crohn’s disease^3,4^. MSCs modulate immune cells through diverse mechanisms including the active production of signaling molecules, release of extracellular vesicles, and signaling via efferocytosis^5–7^. One potent component of this diverse immunomodulatory repertoire is indoleamine-2,3-dioxygenase (IDO), an intracellular enzyme, that has been shown to play a key role in MSC immunomodulatory function. Inhibition of IDO *in vitro* eliminates the suppressive action of MSCs towards activated peripheral blood mononuclear cells (PBMCs)^8^. *In vivo*, infusion of MSCs can induce tolerance to murine renal allografts in an IDO-dependent manner^9^, and overexpression of IDO in MSCs enhances long-term tolerance in a rabbit renal allograft model^10^. As a critical facet of MSCs’ immunomodulatory profile, understanding how MSCs regulate expression and activity of IDO is key to maximizing MSC therapeutic potential.

Expression of IDO in human MSCs is not constitutive, but is induced by cues in the inflammatory environment, predominately interferon gamma (IFNγ)^11^. In contrast to dendritic cells, which can induce IDO in response to a variety of cytokines, MSCs are critically dependent on IFNγ exposure to stimulate IDO expression at both the mRNA and protein level^12^. The effect of IFNγ exposure on IDO protein levels is diminished as MSCs are cultured for extended periods of time and reach senescence^13^. Although senescent MSCs transcribe IDO at a similar rate as earlier passage cells, IDO protein concentration is lower due to proteasomal degradation^14^. Other cytokines, such as tumor necrosis factor alpha (TNFα) and interleukin-1β (IL-1β) augment IDO expression when combined with IFNγ, but are insufficient alone to induce transcription^15,16^. In addition, intact IFNγR signaling and glycolytic metabolism are also both necessary for IFNγ to induce MSC’s immunomodulatory phenotype^17^. Thus, MSCs regulate IDO expression and immunomodulatory phenotype by integrating diverse cues within their environment.

Environmental cues can be manipulated during the biomanufacturing of MSCs to pre-activate an immunomodulatory phenotype, a process called pre-licensing^18^. Pre-licensing has shown benefits across a variety of MSC donors,^18–20^ tissue sources, and settings but the pre-licensing methods used have been highly variable. Pre-licensing with IFNγ doses ranging from 5 to 200 ng/ml and durations ranging from 2 hours (h) to 4 days have been reported^8,21–24^. In addition, Menard *et al.* has shown that after pre-licensing with 10 ng/mL IFNγ and 15 ng/mL TNFα, MSCs isolated from different tissue sources and cultured within different growth medias have differential IDO expression and immunomodulatory potency^25^. Thus, the relative role of the source of MSCs versus the dose and duration of pre-licensing required to enhance MSC immunomodulatory function still needs to be identified to scale biomanufacturing of pre-licensed MSCs (PL-MSCs). Without optimizing these parameters, the best case scenario is an increase in cost without improved performance, while the worst case scenario is the loss of therapeutic efficacy. With differences in potency identified as a function of both the source of MSCs as well as the pre-licensing protocol, how these variables interact are critical to moving PL-MSCs into biomanufacturing systems.

While IDO expression is dictated by diverse, interacting cues, herein we sought to take a systematic approach to identify the contributors that dictate MSC expression and maintenance of IDO. After determining the effectiveness of a variety of pre-licensing strategies to enhance IDO in MSCs, these pre-licensing strategies were tested in human PBMC suppression assays to determine how they translated to immune suppression. Insight into the relative role different parameters play in the production and activity of IDO in MSCs and how this altered IDO profile translates to immunomodulation will inform the design of biomanufacturing protocols for PL-MSCs.

## Materials and Methods

### Reagents

All reagents were purchased from major suppliers unless otherwise stated. IFNγ (PeproTech, 300-02), 98% ≥ L-tryptophan (Sigma-Aldrich, T0254-5G) and L-Kynurenine (Sigma-Aldrich, K862). L-kynurenine was utilized as a 5mM stock concentration, solubilized in MEM-α (Biological Industries, 01-042-1A). Heated kynurenine was generated by heating a 5mM L-kynurenine stock at 37°C for 48 h.

### MSC Source and Culture

Bone marrow derived MSCs (MSC1 (00055) and MSC2 (00082)) were obtained from RoosterBio, while umbilical cord (MSC3 (4477)) MSCs were isolated using a tissue explant method, expanded, and characterized in house, as we previously described^20^. All MSCs met the minimal criteria as defined by the International Society for Cellular and Gene Therapy^26^. Prior to experiments, cryopreserved MSCs were thawed and cultured to 70-90% confluence before splitting and re-plating. MSCs were used at passages P4-P6 for experimentation. All MSCs were grown in MEM-α (Biological Industries, 01-042-1A) supplemented with 15% (v/v) premium grade US Origin-ISIA traceable FBS (VWR, 97068-085), 1% (v/v) Penicillin-Streptomycin (Biological Industries, 03-031-1B), and 1% (v/v) L-Glutamine (Gibco, 25030-081).

### IDO Activity Assay

IDO activity was measuring by quantifying the change in the amount of kynurenine in the media via Ehrlich’s reaction as previously described^27,28^. Briefly, A standard curve of known kynurenine concentrations was constructed using a 1:1 serial dilution ranging from 200μM to 3.125μM. Complete growth media was used for the standard curve to ensure a match between the standard and the experimental samples. 200μL of standard kynurenine concentrations and samples was then added to a 96 well plate followed by 100μL of 30% (w/v) trichloroacetic acid (Sigma-Aldrich, T9159-100G) to each well. The 96 well plate was heated to 52°C for 30 min to convert N-formylkynurenine to kynurenine. After heating, the plate was centrifuged at 1200g for 15 minutes. 150μL of supernatant was then collected, mixed, and an equal volume was placed into 2 adjacent wells of a new 96 well plate. 75μL of Ehrlich’s reagent (0.4% p-dimethylaminobenzaldehyde (Sigma-Aldrich, 156447-25G) in glacial acetic acid) was then added to each well for a final volume of 150μL. Kynurenine reacts with Ehrlich’s reagent to create a yellow color proportional to the concentration of kynurenine. Absorbance at 492 nm was then immediately quantified using a plate-based spectrophotometer (Molecular Devices, 60139412). Absorbance values of the known kynurenine standards were used to construct a standard curve, from which the unknown sample concentrations were interpolated.

### MSC and PBMC Co-culture

PBMCs were obtained via isolation from leukocyte reduction cones provided by the DeGowin Blood Center at the University of Iowa Hospitals and Clinics from healthy de-identified donors. PBMC seeding was fixed at 200K and 20K MSCs were plated to establish a 1:10 MSC:PBMC. A 1:10 ratio in our experience typically results in naïve MSCs suppressing 30-60% of PBMC proliferation, providing ample room to detect either improvement or loss of MSC’s immunomodulatory behavior^13^. RPMI (Biological Industries, 01-100-1A-12) supplemented with 10% (v/v) FBS (VWR, 97068-085), 1% (v/v) Penicillin-Streptomycin (Biological Industries, 03-031-1B), and 1% (v/v) L-Glutamine (Gibco, 25030-081) was used for all co-culture and control wells. MSCs were seeded first into a 24 well plate and allowed to attach for 2 h. During the 2 h MSC attachment period, previously cryo-preserved PBMCs were cultured in RPMI for 1 h. After thawing, PBMCs were stained with CFSE dye at a concentration of 1 μM in PBS for 15 min (Biolegend, 423801) according to the manufacturer’s protocol. Following staining and dye neutralization, PBMCs were mixed at a 1:1 ratio with anti-CD3/anti-CD28 Dynabeads and then added to co-culture wells. PBMCs with or without Dynabead activation were used as positive and negative controls, respectively, and used for gating in each experiment. After 6 days, PBMCs were collected, centrifuged at 500g for 5 min, and then resuspended for analysis on an Accuri C6 flow cytometer.

### IDO Western Blot

Cell lysate was collected in complete RIPA buffer (Santa Cruz Biotechnology, sc-24948A) and centrifuged at 8000g for 10 min. Clarified protein was then denatured with 4X LDS sample buffer (Thermo Fisher, B0007) and 10X Bolt reducing agent (Invitrogen, B0009) followed by heating at 95°C for 2 min. Loading volumes of 10-15 μg of protein per well were calculated using a MicroBCA Protein Assay Kit (Bio Basic, SK3061) according to the manufacturer’s specification. Samples were run through a 4-12% Bis-Tris gradient gel followed by transfer to a PVDF membrane. Primary antibody was added which consisted of 5% (w/v) BSA (Sigma-Aldrich, A9647-10G) and either 1:1000 IDO primary antibody (BioLegend, 122402) or 1:10,000 β-Actin antibody (Thermo Fisher, AM4302). Secondary antibody was then added which consisted of 5% (w/v) milk, and either 1:10,000 anti-rabbit HRP conjugated antibody (Santa Cruz, sc-2004) for IDO or 1:10,000 anti-mouse HRP conjugated antibody (Biolegend, 405306). A WesternBright Quantum mix (Advansta, K-12042-D10) was used for HRP substrate. The membrane was then scanned on a C-DiGit Blot Scanner (LI-COR Biosciences, 3600-00) on high sensitivity. The densitometry of each band was calculated using LI-COR Image Studio software. IDO intensities were normalized to β-Actin intensity for each sample.

### Statistics

The statistical tests used for each experiment are listed in each figure caption. GraphPad Prism v8 was used to conduct statistical analyses.

## Results

### MSC production of IDO is dependent on the dose and duration of IFNγ exposure

A wide variety of doses and durations of IFNγ exposure have been employed to pre-license MSCs; however, testing both of these parameters simultaneously has made it difficult to determine the specific dose and duration dependent effects of IFNγ exposure on MSC IDO protein. To isolate the specific contribution of IFNγ dose on IDO protein, we fixed the duration of IFNγ exposure to 2 days and cultured a single bone-marrow MSC donor (MSC2) with doses ranging from 0 to 100 ng/mL. IDO protein levels were detectable at 1 ng/mL and continued to increase in a dose-dependent manner until reaching a plateau at 50 ng/mL (Fig. 1A).

**Figure 1:**
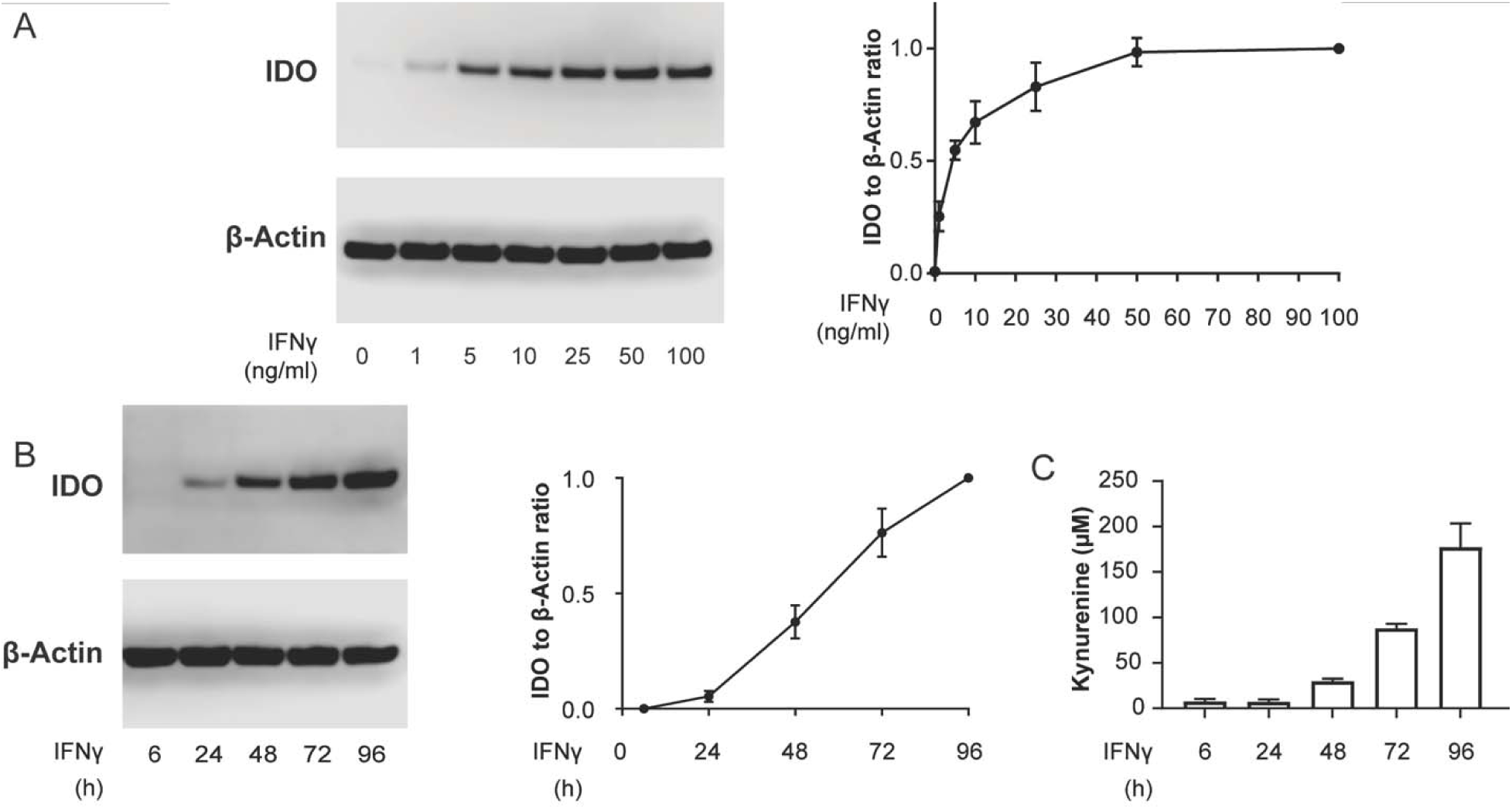
IDO Protein Levels are Dictated by both the Dose and Duration of IFNγ licensing A) Representative western blot and densitometry of MSCs licensed with increasing doses of IFNγ. IDO densitometry was normalized to housekeeping protein β-Actin, and further normalized to the 100 ng/mL IFNγ condition. (N=3 independent experiments, Reporting mean ± SEM). B) Representative western blot and densitometry of MSCs licensed for increasing durations at a fixed 50 ng/mL dose of IFNγ. IDO densitometry was normalized to housekeeping protein β-Actin, and further normalized to the 96 h condition (N=3 independent experiments, Reporting mean ± SEM). C) Activity of IDO, measured as kynurenine concentration at the end of pre-licensing. N=3 independent experiments, paired to Figure 1B. Reporting mean ± SEM.

Next, we sought to determine the contribution of the duration of IFNγ exposure on IDO protein levels and enzymatic activity. We hypothesized that increased duration of IFNγ exposure would lead to enhanced IDO protein levels, as seen with increased IFNγ dose. To test this hypothesis, MSCs were cultured in 50 ng/mL of IFNγ for 6, 24, 72 or 96 h. IDO protein levels increased linearly as a function of the duration of IFNγ exposure with an R^2^ value of 0.899 (p=0.0141, Fig. 1B). Detectable protein levels did not occur until the 24 h mark, however, the minimum time necessary to induce IDO protein may occur anytime within the 6-24 h window. In addition, IDO activity, as measured by the conversion of tryptophan to kynurenine, mirrored what was seen with IDO protein concentration. Kynurenine production was first detectable at 6 h and continued to increase throughout the 96 h period (Fig. 1C), demonstrating that IDO remains active during this extended course of IFNγ exposure.

### Pre-licensing Strategies Increase IDO in all MSCs Relative to Donor Baseline

With larger doses and longer durations of IFNγ exposure shown to enhance IDO protein levels (Fig. 1A,B), we next wanted to determine if these parameters could be utilized as part of a pre-licensing strategy aimed at elevating IDO protein levels in poorer performing donors. Two pre-licensing strategies were tested: a dose based strategy (fixed duration and variable dose) and a duration based strategy (fixed dose and variable duration). MSC immunomodulatory potential and IDO protein concentration have been shown to vary widely between individuals and tissue sources. Therefore, we chose to test these strategies in three independent donors, two bone marrow donors (MSC1, MSC2) and one umbilical cord donor (MSC3).

First, we tested the dose based strategy with either 5 or 50 ng/mL IFNγ exposure for two days. In our previous experiments, 5 and 50 ng/mL of IFNγ exposure induced half and 90% of the maximal IDO protein, respectively; thus, these doses were selected for a mid- and high-level of IDO induction. At the 5 ng/mL dose, MSC3 and MSC2 showed a similar amount of IDO protein, while MSC1 had ∼46% less, indicating a variability at baseline. When treated with high dose of IFNγ, all donors showed a significant increase in IDO protein relative to the low dose (Fig. 2A). In addition, when pre-licensed with a high dose of IFNγ, MSC1 produced roughly equivalent IDO as the other two donors at a low dose. IDO enzymatic activity also increased in all three donors when pre-licensed with high dose IFNγ (Fig. 2B). Therefore, high dose IFNγ during pre-licensing is an effective strategy to increase IDO protein concentration and IDO activity in lower producing donors.

**Figure 2:**
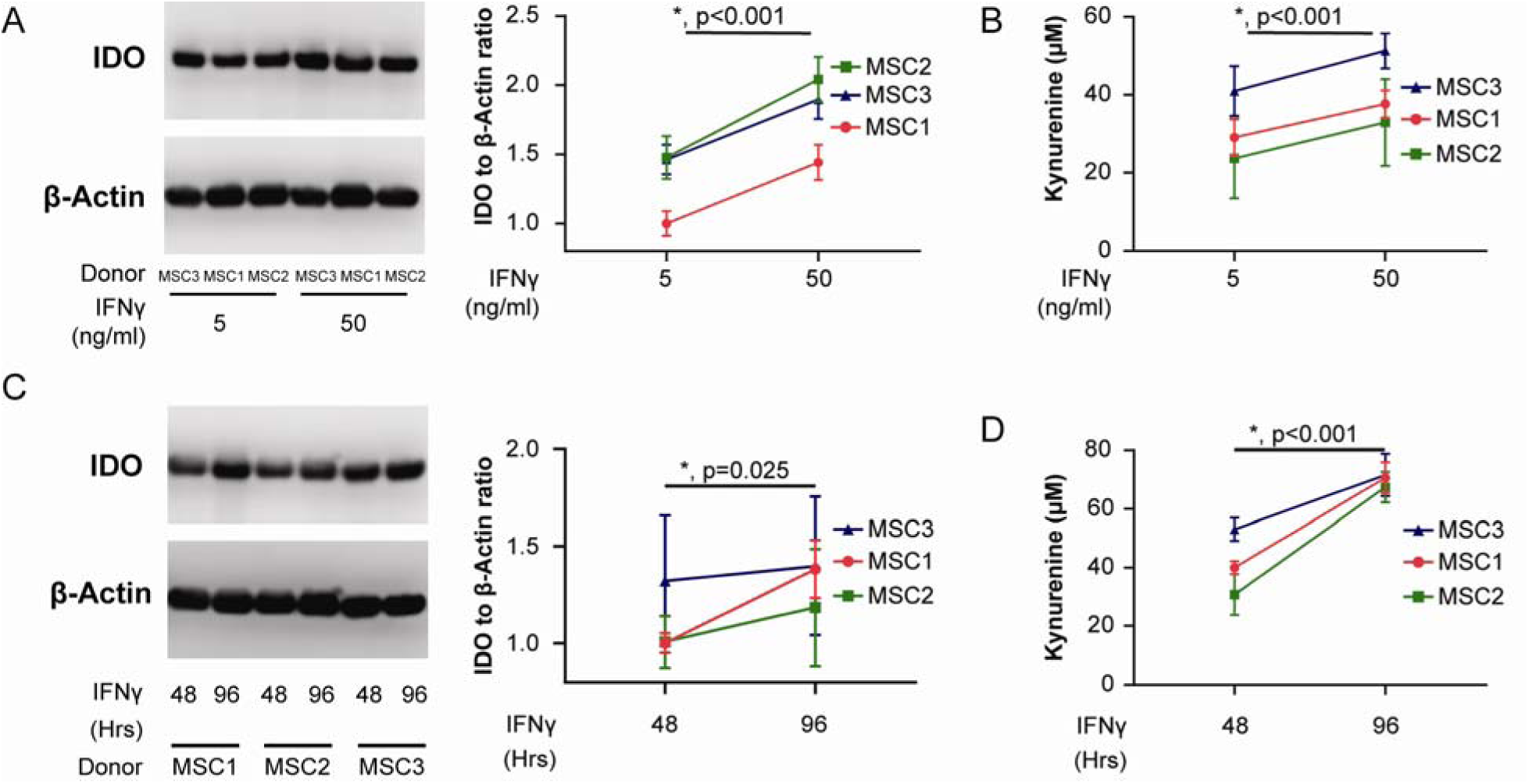
Enhanced Dose and Duration of IFNγ During Pre-licensing is an Effective Strategy to Increase IDO Protein Regardless of Baseline Donor Profile A) Representative western blot and densitometry of MSCs pre-licensed in either 5 or 50 ng/mL IFNγ. IDO densitometry was normalized to housekeeping protein β-Actin, and further normalized to the MSC1, 5 ng/mL IFNγ condition. N=3 independent experiments, mean ± SEM. B) Activity of IDO, measured as kynurenine concentration at the end of pre-licensing. N=3 independent experiments, paired to Figure 1B. Reporting mean ± SEM. C) Representative western blot and densitometry of MSCs pre-licensed for either 48 or 96 h. IDO densitometry was normalized to housekeeping protein β-Actin, and further normalized to the MSC1, 48 h condition. N=3 independent experiments, paired to Figure 2D. Reporting mean ± SEM. D) Activity of IDO, measured as kynurenine concentration at the end of pre-licensing. N=3 independent experiments, paired to Figure 2C. Reporting mean ± SEM. Statistical analysis used for A, B and C was a 2-Way ANOVA with pairing across donors, *p<.05 considered significant.

Next, we sought to determine the effect of duration on IFNγ pre-licensing. To test this, we pre-licensed MSC donors in 10 ng/mL IFNγ for either 48 or 96 h. All donors showed increases in IDO protein when pre-licensed for a longer duration (Fig. 2C). Surprisingly, when comparing the effect of the dose strategy to the duration strategy, the change in IDO protein after longer durations was less than the change observed using the dose strategy (Fig. 2A, C). In addition, the benefit of longer duration varied between donors. Kynurenine concentration increased from 48 to 96 h of pre-licensing across all donors demonstrating that the enzyme remains active throughout the pre-licensing period (Fig. 2D).

### Duration of Pre-licensing Predicts Durability of IDO Protein Post-withdrawal

After determining the individual contribution of the dose and duration of IFNγ during pre-licensing on IDO protein levels and activity in MSCs, we next investigated if the elevated protein induced by a duration-based strategy would persist and remain functional after removal from the pre-licensing environment. Understanding how MSC potency is or is not maintained after removal from a pre-licensing environment is essential for establishing accurate predictions for therapeutic efficacy. To test this, MSCs were pre-licensed in 10 ng/mL IFNγ for two, four, or six days prior to being transferred to a cytokine-free environment for outgrowth (post-withdrawal period) (Fig. 3A). As seen from prior experiments, MSCs pre-licensed for longer durations had elevated IDO protein levels immediately after withdrawal (Fig. 3B). Interestingly, we found that this elevated IDO protein content persisted throughout the 72 h withdrawal period, with the percent of IDO remaining being highest in MSCs pre-licensed for six days (Fig. 3B). In addition, we analyzed kynurenine output at the end of the pre-licensing period, as well as over the course of withdrawal. Immediately after pre-licensing, all three durations resulted in a similar amount of kynurenine output (Fig. 3C). In the post-withdrawal period, kynurenine production was similar 24 h post-withdrawal for all tested durations, however, the output diverged significantly by 48 and 72 h, with MSCs exposed to longer durations of pre-licensing displaying higher levels of kynurenine production (Fig. 3D). Statistical analysis by 2-way ANOVA with Tukey post-hoc test revealed the 6-day pre-licensed MSCs produced more kynurenine compared to the 2 day pre-licensed MSCs at all three time points after withdrawal. Therefore, the duration of IFNγ exposure during pre-licensing increases IDO protein levels leading to enhanced persistence and activity of the protein even after removal from the pre-licensing environment.

**Figure 3:**
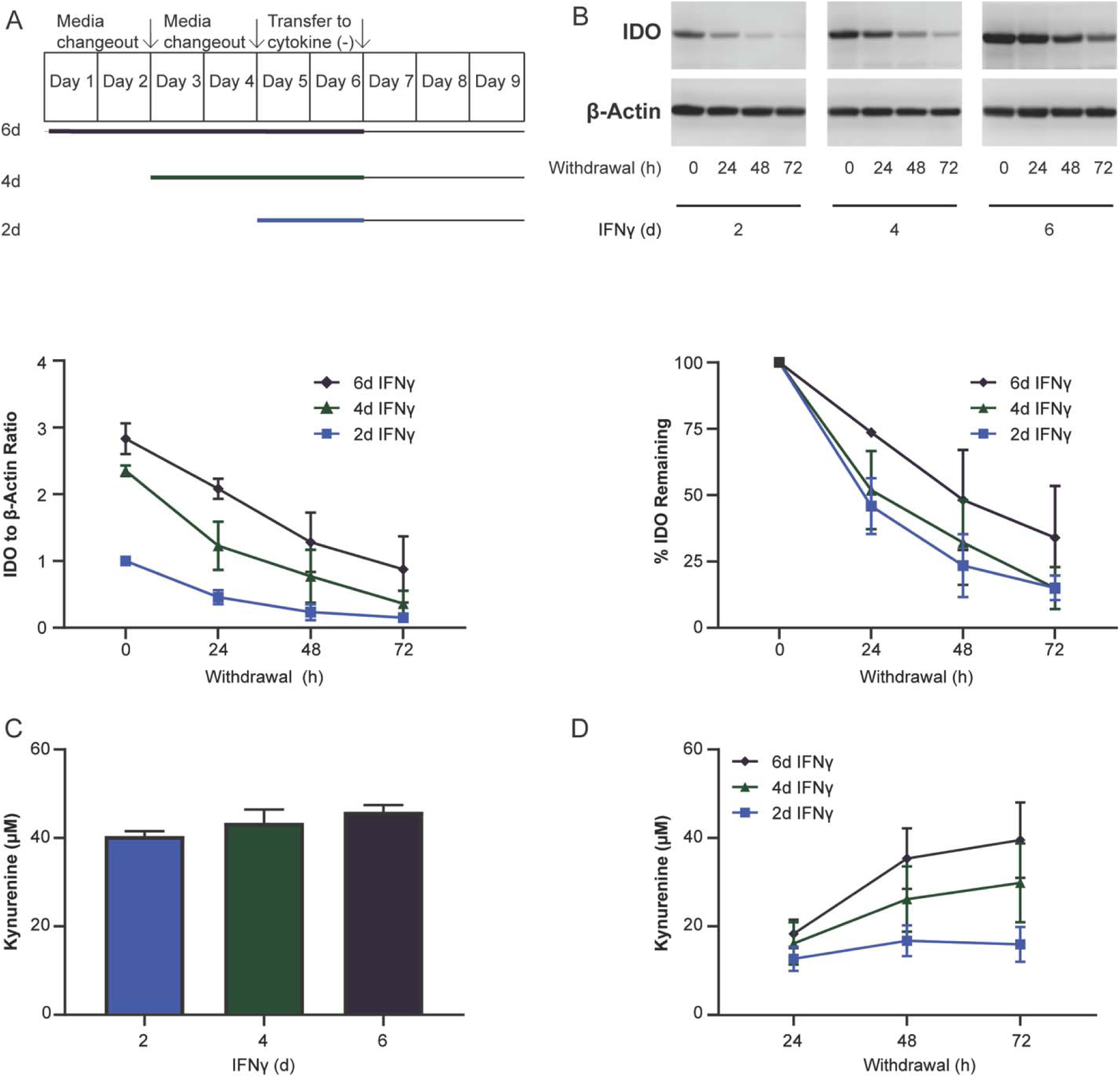
Longer Durations of Pre-licensing Enhances IDO Protein and Activity Post Withdrawal A) Graphical representation of the timing of media changeout, IFNγ stimulation, and withdrawal from cytokines. B) Representative western blot and densitometry of MSCs on the day of withdrawal as well as during the withdrawal period. IDO densitometry was normalized to housekeeping protein β-Actin, and further normalized to the 2 day pre-licensing condition on the day of withdrawal. Percent remaining was calculated as IDO protein for a given pre-licensing duration divided by its IDO protein level on the day of withdrawal. N=3 independent experiments. Reporting mean ± SEM. C) MSCs’ kynurenine concentration for the last 48 h of pre-licensing before withdrawal. N=4 independent experiments. Reporting mean ± SEM. D) Accumulation of kynurenine 24, 48, and 72 h post-withdrawal. N=5 independent experiments. Reporting mean ± SEM.

### Accumulation of Kynurenine Metabolites During Pre-licensing Does Not Affect IDO Protein Concentration or Activity

One concern with pursuing a duration-based strategy of pre-licensing is the potential accumulation of kynurenine and/or kynurenine-derived trace extended aromatic condensation products (TEACOPs). TEACOPs are derivatives of kynurenine that spontaneously form at physiological temperatures and have been shown in other biological systems to have biological activity. Previous studies have identified both of these compounds as regulators of IDO expression in other cell types^29^; however, their effect on MSC regulation of IDO is, as of yet, unexplored. If pre-licensing is to be performed at scale during the biomanufacturing of therapeutic MSCs, it is important to determine the consequence of accumulation of these metabolites on MSCs. To determine the role of kynurenine and spontaneously formed byproducts of kynurenine on IDO within MSCs, MSCs were pre-licensed with 10 ng/mL IFNγ with the addition of either fresh kynurenine or heated-kynurenine (37°C for 48 h) for four days. Unlike other cell types, which show regulation of IDO activity by kynurenine and TEACOPs, neither the addition of fresh nor heated kynurenine had any appreciable impact on the production of IDO protein (Fig. 4A) or IDO enzymatic activity in human MSCs (Fig. 4B). Thus, in MSCs under the conditions examined, kynurenine metabolites do not play a critical role in modulating IDO content or activity.

**Figure 4:**
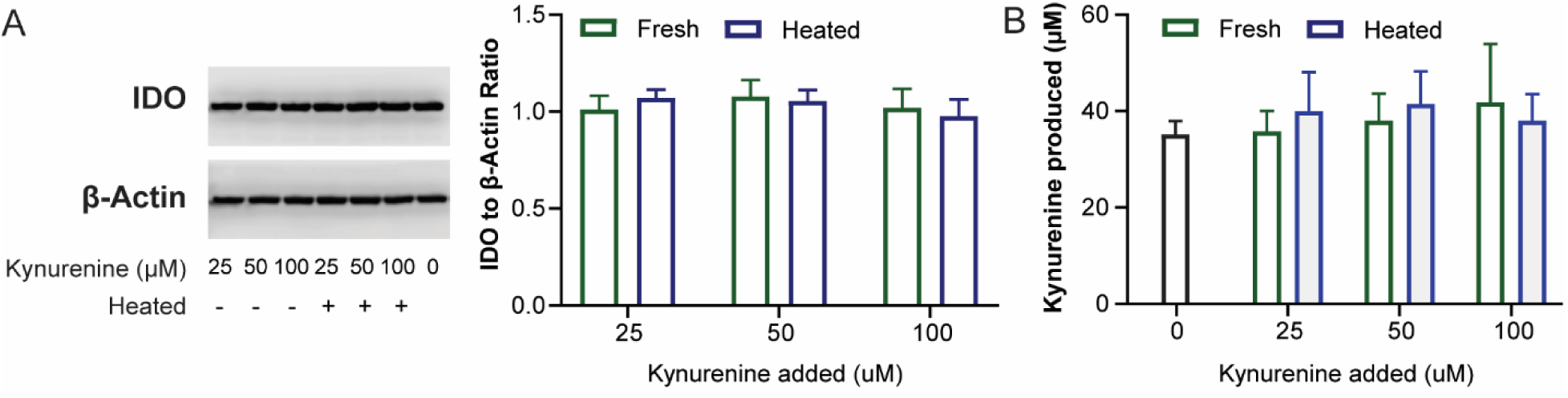
Kynurenine and TEACOPS Do Not Significantly Regulate IDO Protein or Activity A) Representative western blot and quantified densitometry of MSCs pre-licensed with increasing doses of kynurenine with or without heating. IDO protein was normalized to the house keeping gene β-Actin, and further normalized to 0 μM kynurenine condition. N=3 independent experiments. Reporting mean ± SEM. B) Activity of IDO, measured as kynurenine concentration at the end of pre-licensing. Kynurenine produced was calculated by subtracting out signal generated due to kynurenine being added at the beginning of the experiment as measured on an identical plate without MSCs. N=4 independent experiments. Reporting mean ± SEM.

### T Cell Suppression is Enhanced with High Dose IFNγ but Diminished with Longer Pre-licensing

Having isolated the individual contribution of IFNγ dose and duration on IDO protein and activity, we next sought to determine if this translated to enhanced potency in a human PBMC suppression assay. To quantify the immunosuppressive ability of PL-MSCs, we pre-licensed our three donors (MSC1, MSC2, MSC3) using three unique strategies prior to direct contact co-culture with CD3/CD28 activated PBMCs, as well as a naive control, to determine baseline immunosuppressive ability for each donor. Three pre-licensing strategies used were: base pre-licensing (10 ng/mL IFNγ for two days), high dose pre-licensing (50 ng/mL IFNγ for two days), and prolonged duration pre-licensing (10 ng/mL IFNγ for four days). Based on the results of our IDO protein and activity assays, we hypothesized that both high dose and longer duration would improve immunosuppressive potency compared to the base protocol.

Our positive control, base PL-MSCs, showed improved immunosuppressive potency compared to naive MSCs, indicating a benefit to pre-licensing as has previously been shown by us and others^20,23,25,30^,. Additionally, consistent with our hypothesis, all donors benefited from high dose pre-licensing, showing the highest suppression of proliferation across all tested strategies (Fig. 5A, B). Surprisingly, while IDO assays showed prolonged duration results in improved IDO levels and activity, it did not improve suppression of PBMCs compared to naive MSCs and was significantly worse than base pre-licensing (Fig. 5A, B). When comparing base pre-licensing to high-dose pre-licensing, although there was a significant difference between these strategies (p=0.03), the difference was modest (4.9% improvement). Using a 2-way ANOVA to determine the amount of variance accounted for by donor choice versus pre-licensing strategy, most of the variance in immunomodulatory performance was attributable to donor choice (51% of variation) while pre-licensing strategy accounted for 31% of the variance and interactions between MSC donor and pre-licensing strategy accounted for 15% of the variance. Therefore, both the pre-licensing strategy and donor choice have significant roles in determining MSC immune suppression.

**Figure 5:**
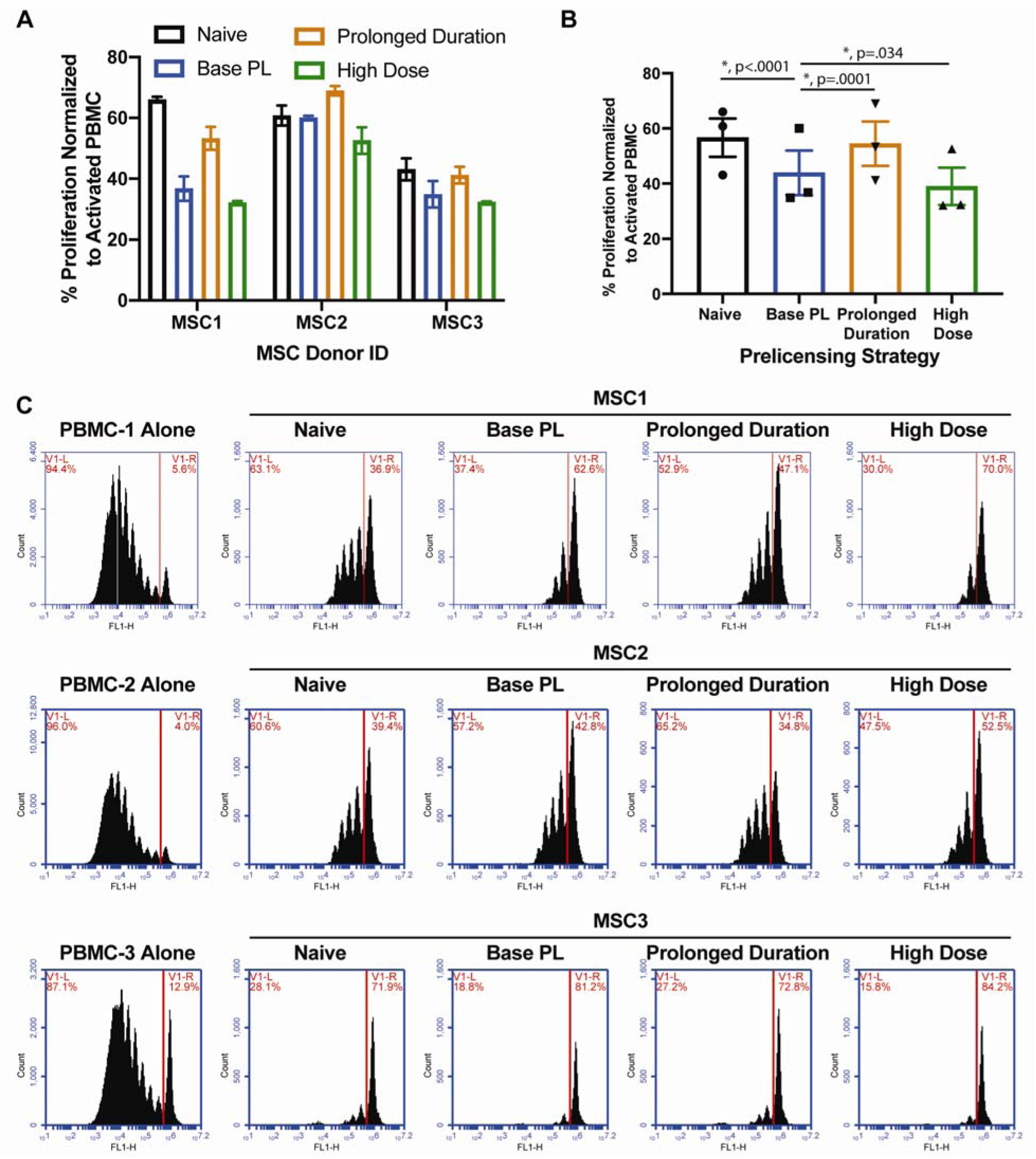
Dose, Duration and Donor are Regulators of MSC Immune Modulation A) Proliferation of PBMCs co-cultured with MSC donors MSC1, MSC2 and MSC3 at a 1:10 PBMC to MSC ratio. %Proliferation was calculated by normalizing to stimulated PBMC control within each experiment(positive control). Reporting Mean ± SD of 3 replicates for each donor. B) Proliferation data from A pooled and replotted to compare the effects of pre-licensing strategies independent of donor. Each bar is a pre-licensing strategy and each data point is the mean %Proliferation for an MSC donor in that strategy. N=3 independent donors, Reporting Mean ± SEM, 2-Way ANOVA with Dunnett’s multiple comparisons test, *p<.05 considered significant. C) Representative flow plots of CFSE-stained PBMCs upon co-culture with each MSC donor under each pre-licensing condition. Red Line denotes gate used to calculate %PBMC proliferation reported in A, B.

The differences in donor potency varied widely without pre-licensing (Naïve, Fig. 5A). Interestingly, while the trends between pre-licensing strategies were maintained across donors, the magnitude of this benefit varied substantially across donors with MSC1 benefiting the most, MSC3 receiving an intermediate benefit, and MSC2 benefiting the least (Fig. 5A). Regardless of how pre-licensing affected the different donor’s potency, MSC3 was more suppressive than either MSC1 or MSC2. Our data shows that the effect of pre-licensing is not uniform, but the product of an interaction between the pre-licensing strategy and the intrinsic properties of the MSC donor.

## Discussion

While MSCs’ immunomodulatory properties have been extensively documented, these properties are often assumed to be innate characteristics rather than dynamic cell states. Here we focused on MSC regulation of IDO, a critical immunoregulatory enzyme shown to correlate with MSC suppression of PBMCs^31^, and systematically separated the role of the dose of IFNγ stimulation, the duration of pre-licensing, and the MSC donor. These studies were motivated both by a desire to uncover the dynamics of MSC activation into an immunomodulatory state as well as to identify parameters that need to be tuned to biomanufacture pre-licensed MSCs at a therapeutic scale. As expected, we found MSCs increase in IDO content to be dependent on both the dose and duration of IFNγ pre-licensing (Fig. 1 A, B). Interestingly, increasing the dose of IFNγ had a more consistent effect across MSC donors than increasing the duration of pre-licensing. Increasing the dose of IFNγ from 5 to 50 ng/mL led to an increase in IDO protein and production of kynurenine to a very similar degree across all donors tested (Fig. 2A, B). In contrast, the benefit of a duration based strategy was found to be more donor dependent (Fig. 2C). Thus, at the time of withdrawal, a high-dose pre-licensing strategy appears to favor higher levels of IDO compared to an extended duration strategy.

Next, we wished to determine how pre-licensing affected the durability of IDO. While identification of pre-licensing protocols that maximize IDO activity are desirable, they are only useful if IDO remains functional once the cells are transplanted. If IDO is degraded immediately after removal, pre-licensing is of little use. We hypothesized that increased duration of pre-licensing would enhance IDO protein levels, not only on the day of withdrawal, but also throughout the days following withdrawal. Both of our predictions, that IDO protein would be elevated on the day of withdrawal as well as on subsequent days after removal of IFNγ, were supported by the data (Fig. 3B). In addition, IDO enzymatic activity was maintained longer after withdrawal for MSCs that were pre-licensed for longer durations (Fig. 3D). Thus, prolonged pre-licensing appears beneficial both for the levels of IDO protein and the longevity of IDO after removal from the pre-licensing environment.

Next, we examined the role of kynurenines on MSC pre-licensing. Both kynurenine and kynurenine derivatives (TEACOPs) are known ligands for the aryl hydrocarbon receptor (AhR), which has been shown to regulate IDO expression in other cell types^29^. Thus, depending on the type of regulation, accumulation of such byproducts could be either beneficial or detrimental to the manufacturing of potent PL-MSCs. Seok et. al. showed that TEACOPs spontaneously form at physiological temperatures and are potent ligands for AhR. Once bound, the AhR-ligand complex translocates to the nucleus and regulates IDO transcription^32^. AhR regulation can either augment or inhibit transcription of target genes and is dependent upon cell type and the ligand that binds AhR^32^. However, the role of AhR and AhR ligands on MSCs has not been explored. While Seok et al. showed TEACOPS are potent AhR activators in COS-1 cells from African green monkey, exposure of human MSCs to kynurenine or kynurenine that had been heated to 37°C for 4 days did not have an appreciable effect on IDO protein or activity in human MSCs (Fig. 4A, B). Thus, selective removal or addition of these compounds during biomanufacturing does not appear to be a strategy that would augment IDO protein concentration or activity in PL-MSCs.

With the effect of IFNγ dose and duration of pre-licensing on IDO protein and activity tested, we next determined how these different pre-licensing strategies translate to MSC immune suppression. Based on our observations of MSC IDO, we expected both the high-dose strategy and the extended-duration strategy to lead to a significant increase in MSC mediated suppression of PBMCs. To our surprise, enhancing the dose of IFNγ five-fold only moderately increased the level of PBMC suppression (Fig. 5A, B). Even more surprising was that extending the duration of pre-licensing actually had a detrimental effect on MSC potency (Fig. 5A, B). Thus, we found that MSCs immunomodulatory phenotype cannot be captured using a single surrogate marker, in this case IDO, highlighting the need for multiple metrics to be used to predict MSC function^33^. In addition, while we initially thought the dose and duration of pre-licensing would be the most critical parameters to optimize, we found the donor itself had the largest influence on the potency of PL-MSCs. While base and high-dose pre-licensing had a consistent effect across donors, the donor of MSCs itself was responsible for most of the variance in immune suppression observed, highlighting the critical role of the starting cellular material (Fig. 5A). Pre-licensing itself cannot be used to overcome inherent donor impotency and screening of donors for allogeneic use is necessary to maximize the immunomodulatory benefit of pre-licensing.

The strengths of this study lie in our methodical approach to separate the influence of dose, duration, and donor on MSC regulation of IDO and the analysis of MSC immunomodulatory potency using three surrogate markers: IDO protein, IDO activity, and PBMC suppression. However, several limitations should be noted. As the donor source was the largest contributor to MSC immunomodulatory function, MSCs collected from different donors, tissues, and cell culture systems may require a different ‘optimal’ dose of IFNγ and duration of pre-licensing to maximize their therapeutic potency. The strategy outlined here can thus, serve as a template for analyzing a range of doses and durations for specific lots of MSCs. Second, all experiments in this study were performed in a classic MSC media formulation (MEM-α, 15% FBS, 1% L-Glutamine, 1% Penicillin-Streptomycin). We used the highest quality US Origin ISIA certified FBS available, This was done to make our work relevant to both the large body of MSC literature using FBS as well as current MSC products that are being tested in clinical studies^34^. MSCs grown in alternate media formulations could generate MSCs that respond distinctly to cytokine stimulation^35,36^.

## Conclusion

The source of MSCs and the process used to manufacture them prior to transplantation have a major influence on the immunosuppressive properties of MSCs. MSCs dynamically upregulate IDO in an IFNγ dose dependent manner and continue to upregulate IDO during extended periods of IFNγ exposure. However, despite elevated levels of IDO and maintained IDO activity, prolonged exposure to IFNγ can lead to MSCs immunosuppressive potency reverting to levels similar to naïve MSCs. In addition, while both dose and duration can be used to modulate MSC production of IDO, the MSC donor played the biggest role in determining MSCs’ immunosuppressive potency. Thus, the dose and duration of pre-licensing should be tailored to optimize performance of specific donors and an emphasis on donor selection is needed to realize significant benefits of pre-licensing.

## Data Availability

The data used to support the findings of this study are available from the corresponding author upon request.

